# Tandem repeat variation in human and great ape populations and its impact on gene expression divergence

**DOI:** 10.1101/015784

**Authors:** Tugce Bilgin Sonay, Tiago Carvalho, Mark D. Robinson, Maja P. Greminger, Michael Krützen, David Comas, Gareth Highnam, David Mittelman, Andrew Sharp, Tomàs-Marques Bonet, Andreas Wagner

**Author notes:** equal contribution (Tugce Bilgin Sonay, Tiago Carvalho, Tomàs-Marques Bonet, Andreas Wagner). Corresponding authors, address: University of Zurich, Institute of Evolutionary Biology and Environmental Sciences, Building Y27-J-50, Winterthurerstrasse 190, CH-8057, Zürich, Switzerland. address: Institute of Evolutionary Biology (CSIC-Universitat Pompeu Fabra), Dr. Aiguader 88, 08003 Barcelona, Spain.

## Abstract

Tandem repeats (TR) are stretches of DNA that are highly variable in length and mutate rapidly, and thus an important source of genetic variation. This variation is highly informative for population and conservation genetics, and has also been associated with several pathological conditions and with gene expression regulation. However, genome-wide surveys of TR variation have been scarce due to the technical difficulties derived from short-read technology.

Here, we explored the genome-wide diversity of TRs in a panel of 83 human and nonhuman great ape genomes, and their impact on gene expression evolution. We found that populations and species diversity patterns can be efficiently captured with short TRs (repeat unit length 1-5 base pairs) with potential applications in conservation genetics. We also examined the potential evolutionary role of TRs in gene expression differences between humans and primates by using 30,275 larger TRs (repeat unit length 2-50 base pairs). About one third of the 13,035 one-to-one orthologous genes contained TRs within 5 kilobase pairs of their transcription start site, and had higher expression divergence than genes without such TRs. The same observation held for genes with repeats in their 3’ untranslated region, in introns, and in exons. Using our polymorphism data for the shortest TRs, we found that genes with polymorphic repeats in their promoters showed higher expression divergence in humans and chimpanzees compared to genes with fixed or no TRs in the promoters. Our findings highlight the potential contribution of TRs to recent human evolution through gene regulation.

## INTRODUCTION

Tandem Repeats (TRs) are DNA tracts in which a short base-pair motif, the repeat unit, is repeated several times in tandem, i.e., in a closely spaced, head-to-tail orientation. They are among the most variable loci, experiencing mutations in the number of repeat units that are 100 to 100,000 times more frequent than point mutations, and occur at a rate of 10^-3^ to 10^-7^ copy number alterations per cell division (Weber and Wong 1993; Brinkmann et al. 1998; Li et al. 2002; Legendre et al. 2007).

Due to their unique properties, TRs have been extensively used as molecular markers in many population genetic studies (Ellegren 2004). Past technical constraints, however, limited the number of microsatellites that could be easily genotyped. For this reason, most TR-based studies of human diversity and interspecies genetic divergence, were greatly restricted (Rosenberg et al. 2002; Molla et al. 2009; Pemberton et al. 2009; Tishkoff et al. 2009; Sun et al. 2012; Pemberton et al. 2013), or focused on comparing reference genomes (Webster et al. 2002; Kelkar et al. 2008; Payseur et al. 2011; Kelkar et al. 2011; Loire et al. 2013). However, recent advances in sequencing methodology (reviewed in Mardis 2008 and Metzker 2010), and the development of novel software that can systematically genotype repeats at a genome-wide scale (reviewed in Lim et al. 2013), has permitted analysis of several thousand loci from multiple individuals in a cost-effective manner (McIver et al. 2011, 2013; Willems et al. 2014). Furthermore, other recently developed methods which specifically target repeats also show great promise, and will be able to improve both the reliability and scope of repeat landscape analysis (Guilmatre et al. 2013; Duitama et al. 2014; Carlson et al. 2014).

In eukaryotes, TRs located in coding regions tend to occur in genes associated with transcriptional regulation, DNA binding, protein–protein binding, and developmental processes (Vinces et al. 2009; Gemayel et al. 2010). These consistent functional enrichments suggest important functional roles for TRs. In fact, TRs are emerging as good candidates for a type of genomic variation that can directly alter gene expression (Rockman and Wray 2002; Kashi and King 2006; Vinces et al. 2009; Gemayel et al. 2010). Because gene expression changes contribute to the fundamental differences between humans and other species (King and Wilson 1975), it is imperative to study mechanisms that may permit rapid expression changes on short evolutionary time scales (Wray et al. 2003; Tirosh et al. 2006; Landry et al. 2007; Choi and Kim 2008; Tirosh et al. 2009). Promoter features such as TATA boxes, nucleosome density, as well as tracts of TRs can mediate such changes (Tirosh et al. 2009). Thus, the high incidence of TR in regulatory regions (Gemayel et al. 2010; Payseur et al. 2011) and their high mutability (Weber and Wong 1993; Brinkmann et al. 1998; Li et al. 2002; Legendre et al. 2007), suggests that it may be important to study TR variation to understand fundamental differences in gene expression across species and populations. In particular, since TRs constitute 3% of the human genome (Lander et al. 2001) and are dramatically enriched in promoter regions (Vinces et al. 2009; Sawaya et al. 2013) clarifying their functional role may provide important insights for the human biology field.

Much of our knowledge of nonhuman great apes diversity stems from studies which typically focused on variation of a few dozen microsatellite loci (Reinartz et al. 2000; Warren et al. 2000; Zhang et al. 2001; Goossens et al. 2006; Becquet et al. 2007; Bergl and Vigilant 2007; Arora et al. 2010; Wegmann and Excoffier 2010; Gonder et al. 2011; Nater et al. 2013; Fünfstück et al. 2014). This view has been recently complemented with a range of publications including now a comprehensive catalog of great ape diversity using SNP datasets (Locke et al. 2011; Vallender 2011; Prado-Martinez et al. 2013; Scally et al. 2013; Greminger et al. 2014; McManus et al. 2014).

Because different marker types are characterized by different evolutionary rates and modes, they can give us complementary insights into the evolutionary history and present diversity patterns within and between closely related species. For this reason, analyses of the full spectrum of genomic variation in human and nonhuman great ape populations are critical, and in that regard have already proved very informative (Prado-Martinez et al. 2013; Sudmant et al. 2013; Hormozdiari et al. 2013). In light of these facts, a more complete description of the repeat landscape in great apes can help us further understand what makes us human, and how human populations have been shaped by processes such as natural selection and demographic history. This information will also be valuable for conservation efforts of nonhuman primates, since proper population management is greatly enhanced by the availability of molecular markers that allow for efficient diversity assessment from, for example, non-invasive samples.

To characterize the extent of TR variation in humans and great apes, as well as its influence on patterns of gene expression, we examined both TR polymorphisms and divergence in different human populations and their closest primate relatives, chimpanzees and other great apes. We also investigated a potential expression divergence between human and chimpanzees based on non-randomly associated TRs located at promoters.

## RESULTS

### Divergence and Population diversity of TRs

We first identified patterns of TR birth/death across human and nonhuman great apes (chimpanzees, gorillas and orangutans), considering TRs whose repeat units ranged from 1 to 5 base pairs in length, and with a total repeat length of at least seven base pairs, if the repeat unit was one base pair long, and six base pairs otherwise. To this end, we used Tandem Repeat Finder (TRF) (Benson 1999) to identify TR candidate loci in the reference genomes of these taxa and looked for TRs with the same repeat motif in the corresponding regions in the reference genomes of the other three taxa (see methods). We found that more than 70% of the repeat loci annotated in the human genome are shared with chimpanzees, gorillas and orangutans. Furthermore, the degree of repeat conservation somewhat reflects the evolutionary history of these species, with human and chimpanzee sharing the most repeat loci (86.6–87.9%), and orangutan sharing the fewest repeat loci with the other three taxa (68.5%-73%) (Supplementary Table 1). We also observed that very few repeat loci are shared exclusively between any two taxa (0.9-5.1%), and that the percentage of repeat loci specific to each taxon is highest in orangutans (21.8%), followed by gorillas (9.9%), chimpanzees (7.1%), and finally humans (6.2%) (Supplementary Table 2). We performed a gene ontology (GO) enrichment analysis on the genes whose promoters contained repeat loci in the human reference genome, but whose corresponding regions in other taxa lacked a repeat, and found that these genes relate to a wide range of biological processes, such as cell communication and signaling (Supplementary Table 3).

To assess the degree of polymorphism of these repeats in each species, we analyzed the genotype variation in a total of 6,965,726 TR loci in humans, 4,006,024 in chimpanzees and bonobos, 3,815,198 in gorillas, and 2,313,198 in orangutans. Specifically, these were genotyped in a panel of 27 humans (2 from Europe, 2 from Asia, 1 from America, 1 from Oceania and 21 sampled throughout Africa, of which 9 are newly sequenced), 16 chimpanzees (4 subspecies) and 12 bonobos, 18 gorillas (2 species, 3 subspecies), and 10 orangutans (2 species) respectively. To produce this repeat catalog we used the human reference genome genome assembly (NCBI 37, hg19) as a reference for all individuals, in order to facilitate coordinate comparison and to take advantage of the thorough human gene annotation. Additionally, to avoid potential biases resulting from a single reference sequence, we also genotyped the corresponding TRs in the reference genome of each genus and filtered out any repeats whose genotypes were not concordant when comparing the results using both references (see Methods).

For the genotyped repeat loci, we estimated average differences in absolute repeat number relative to those annotated on the hg19 human reference genome in different subspecies and species of nonhuman primate taxa, and in three human groups (Non-Africans, African hunter-gatherers and all other Africans). The total amount of repeat number differences accumulated by each group was consistent with what was expected based on their phylogenetic distance to the human reference genome. Specifically, orangutans showed the most repeat number differences, followed by gorillas, bonobos, chimpanzees, and finally humans, where African hunter-gatherers displayed slightly more differences relative than other Africans and Non-Africans (Figure 1A). We also found that the general patterns of heterozygosity between species and subspecies were highly concordant with those of SNP diversity in great apes on a similar dataset (Prado-Martinez et al. 2013)(Figure 1B).

**Figure 1.**
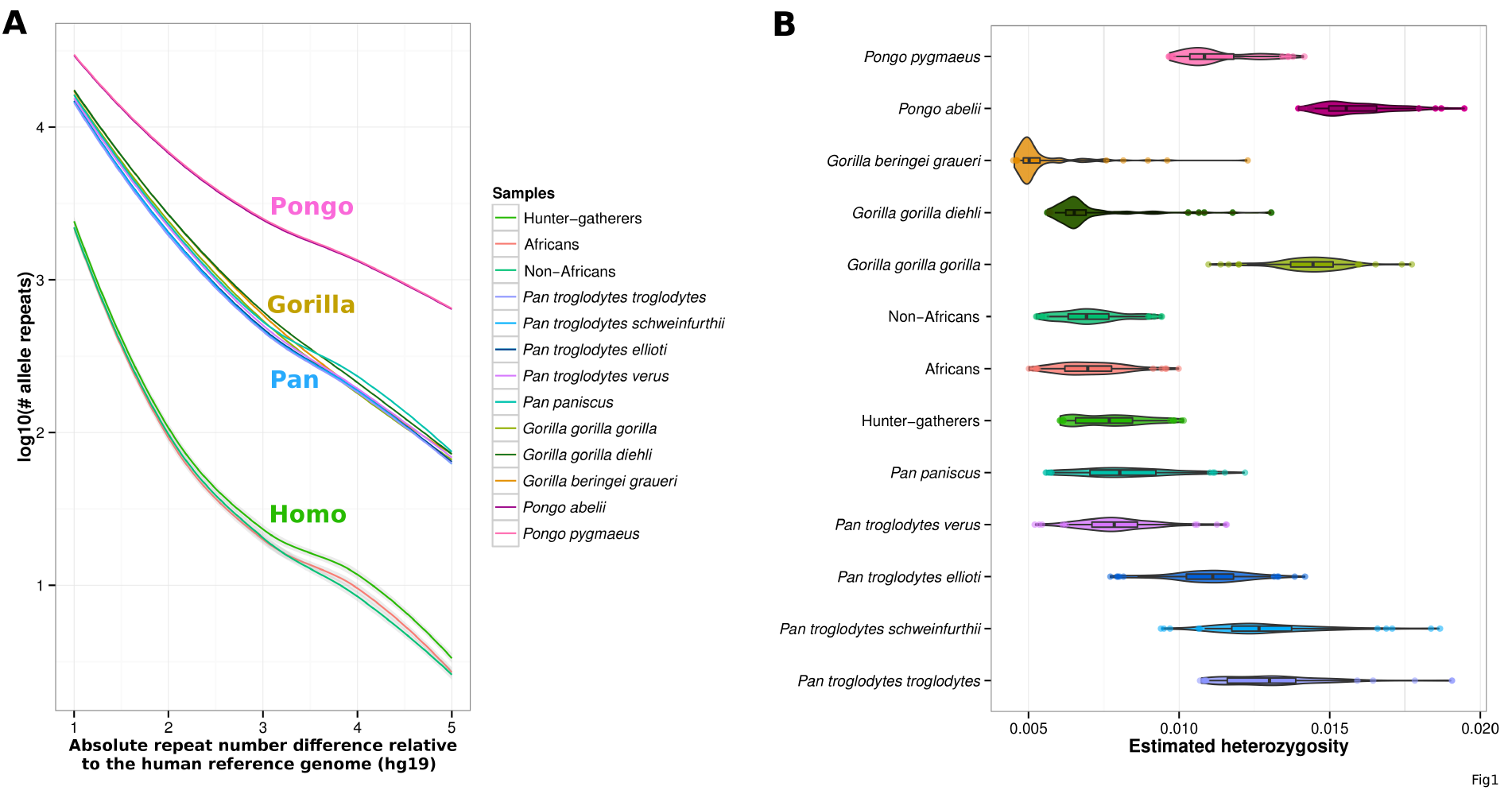
Repeat copy number differences relative to human reference genome (hg19) and heterozygosity estimates for several nonhuman great apes species and human groups. A) Absolute repeat number differences relative to the human reference genome estimated to occur for different human groups and subspecies/species of nonhuman primate taxa. The x axis shows the number of repeat copy number differences and the y axis the number of events for each repeat number difference in log10 scale. As expected, humans show the fewest differences relative to the human reference genome, and are followed by chimpanzees, bonobos, gorillas and orangutans. Within humans, African hunter-gatherers show the most variation, followed by Africans and non-Africans. B) Heterozygosity estimates for different human groups and subspecies/species of nonhuman primate taxa. These results show great concordance with previous genetic surveys using millions of SNPs.

In order to summarize the genetic diversity information contained in the TRs, we computed their allele frequencies within human and non-human great ape populations, and performed Principal Components Analysis (PCA) individually for each taxon. We found that for humans, known diversity patterns among human populations can be accurately captured. Specifically, the first principal component explained up to 10% of the variance, clearly distinguishing a group encompassing North Africans and Non-African individuals from the rest of African samples (Supplementary Figure S1A). In addition, some African samples, such as the South African Bantu, or the North Chad, which are respectively geographically closer to African hunter-gatherer and to North-African populations, are also closer to each other in the PCA. The second and third principal components separately distinguish each of the two African hunter-gatherer populations, San and Mbuti, previously reported to be among two of the most genetically diverse human populations (Tishkoff et al. 2009), from other African and Non-African populations.

Using the PCA approach, we were able to partition not only between great ape subspecies, but also between different great ape populations according to their geographical origin, with just up to four principal components. Even for the great apes with a very complex taxonomy (chimpanzees), each of the first three principal components, respectively, separated the Nigeria-Cameroon and West African chimpanzees from Central and Eastern chimpanzees (19% explained variance), followed by Nigeria-Cameroon and West African chimpanzees (13% explained variance), and finally Central and Eastern chimpanzees (10% explained variance) (Figure 2A). In addition, the fourth principal component identified substructure in the central and eastern subspecies, which correspond to the sample's geographical origin. Likewise, for gorillas and orangutans the PCA could clearly separate between species and subspecies (Supplementary Figure S1C and S1E). These same population structure patterns had been previously observed (Prado-Martinez et al. 2013) using millions of SNPs, and might be best explained by the widespread geographical distribution of chimpanzee and gorilla species and subspecies across equatorial Africa, as well as across and within the Borneo and Sumatra islands for orangutans.

**Figure 2.**
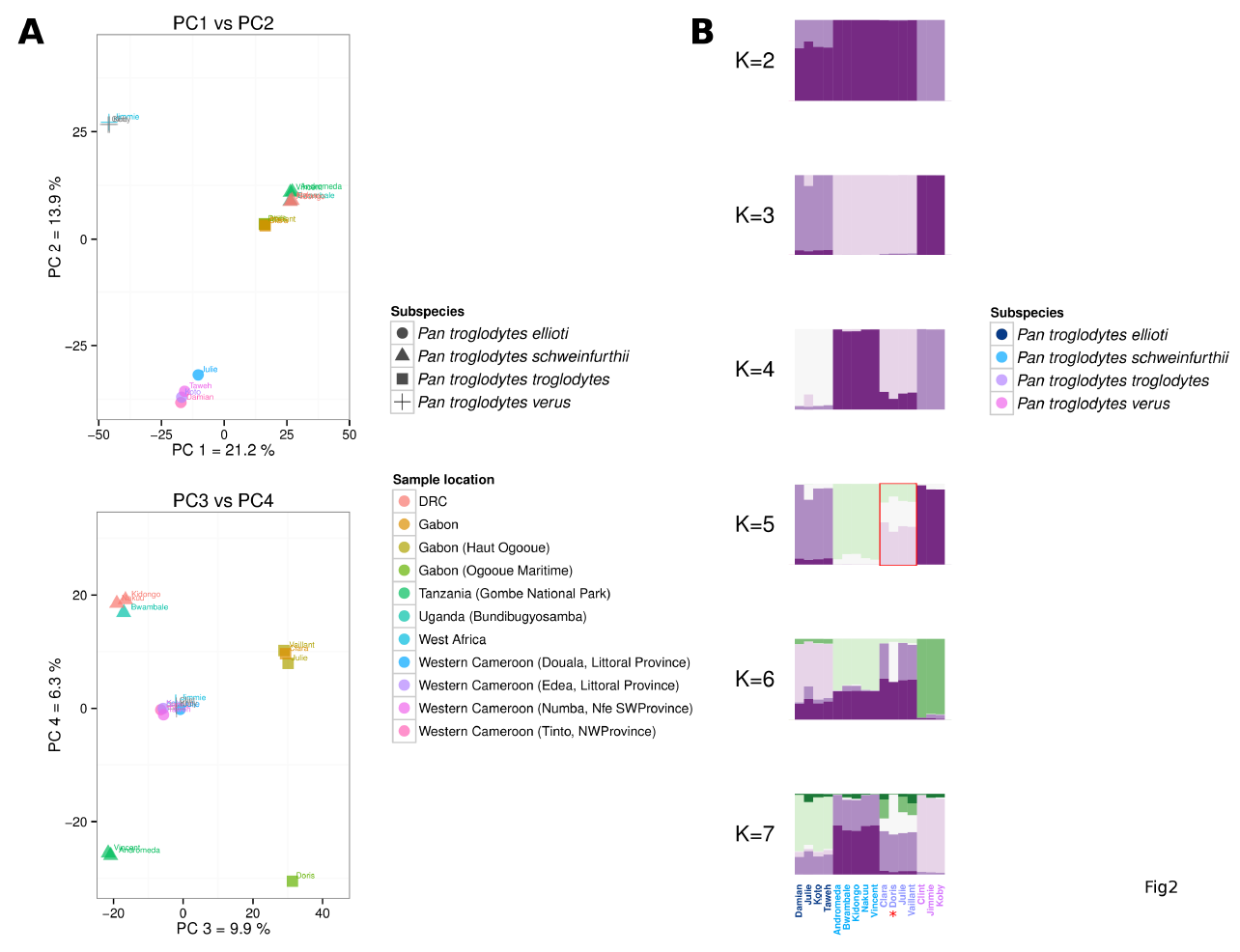
Population structure analysis of chimpanzees, using a principal component analysis (PCA) and STRUCTURE. A) In the PCA plots each axis corresponds to a principal component, and the percentage of the total variation it explains. The upper plot shows the two first components, and the bottom plot the third and the fourth principal components (PC). Shapes correspond to different chimpanzee subspecies, and colours indicate geographical origin (DRC stands for Democratic Republic of the Congo). The first three principal components separate the four chimpanzee subspecies currently described in the literature, and the fourth highlights within-subspecies structure due to the geographical origin of the samples. B) Each barplot shows the results obtained with STRUCTURE when considering different K populations (shown left of the plots) into which the samples (each represented by a bar) were partitioned into. The structure patterns observed are shown to correspond to subspecies designation (K=2 through K=4). The number K of populations best supported by the data is K=5, according to Evanno's method. This corresponds to further substructure within the central chimpanzee population (red box), which may be due to genetic differences between samples belonging to populations from eastern and western Gabon, also observed in the PCA. Specifically, Doris (indicated by a red asterisk), which originates from western Gabon possesses a different genetic component from the eastern central chimpanzee samples.

We obtained similar results when employing STRUCTURE (Pritchard et al. 2000) on the same set of markers used for the PCA for each taxon (Supplementary Figure S2). We then used these observations to assess what number of clusters best explains the substructure present within each taxon by calculating Evanno's delta K (Evanno et al. 2005). With exception of common chimpanzees, for which the optimal number of clusters identified was five and thus one more than the number of subspecies currently reported (Bowden et al. 2012), our observations on all other taxa, for which species and subspecies are described, underscore their common taxonomical designations. The additional cluster in chimpanzees highlights deep population structure in chimpanzees, probably due to geographical substructure that exists within the central chimpanzees population, as suggested by visual inspection of the STRUCTURE plots (Figure 2B).

These observations are directly applicable for conservation genetics. Since TRs are the standard molecular marker in the field, we compiled a list of ancestry-informative markers (AIMs) for all taxa (chimpanzees, gorillas, and orangutans). To do this we searched for repeat loci in species and subspecies where at least 75% of the samples had been genotyped and selected those in which all alleles present were specific to a particular group (Supplementary Table 4). We found a total of 3,522 loci that can now be used to differentiate between subspecies.

To get a sense of the amount of population differentiation present within each taxon, we calculated the standardized Rst value (Slatkin 1995) between all pairs of individuals. When visualized on a heat map in which samples are arranged according to their taxonomical or population designation, clusters of low pairwise Rst values can be observed, highlighting the higher degree of genetic similarity between samples from the same species, subspecies or even population. These are particularly noticeable for nonhuman great apes, and to a lesser extent in humans, which is to be expected since humans are genetically less differentiated than other great apes (Enard and Paabo 2004; Marques-Bonet et al. 2009)(Figure 3).

**Figure 3.**
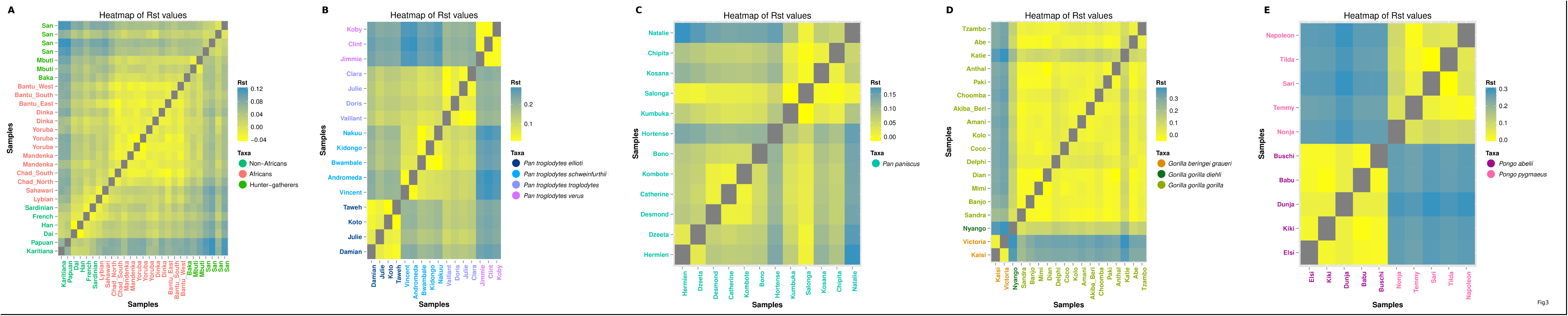
Heatmap plots with Rst values for all great ape taxa. Heatmap plots of Rst for human and nonhuman primate taxa, produced by comparing samples in a pairwise fashion. Yellow indicates a low Rst value, and corresponds to high similarity between a pair of samples, while blue indicates high Rst and dissimilar samples. Clusters of higher similarity between samples corresponding to populations, subspecies, and species can be observed.

### The impact of TR on gene expression

To identify all genes with any kind of TR in the 5 kbps upstream from the transcription start site (TSS), we used a set of 13,035 one-to-one orthologous genes in the reference genome assemblies of humans (NCBI GRCh37.p10), chimpanzees (CHIMP2.1.4), and macaques (MMUL_1) (see Methods). We found that on average 29–31% of these genes harbored TRs (3,820, 3,910 and 4,032 for human, chimpanzee and macaque, respectively). To check the functionality of these repeats, we carried out a randomization test (n=500) using data from ENCODE (Material et al. 2004) on DNase hypersensitive site locations human lymphoblastoid cells (Sabo et al. 2006) which are accessible regions of DNA, associated with gene regulatory elements (Gross and Garrard 1988). We found that 60% of TRs overlap with a DNase hypersensitive site (P-value = 10^-350^). The significant enrichment of repeats in DNase hypersensitive sites suggests that a substantial part of repeat sequences could potentially be involved in gene regulation. Based on this premise, we used previously obtained RNA-seq gene expression data (Brawand et al. 2011) to assess whether genes that contain TRs in their promoters have higher expression divergence compared to genes without repeats in their promoter region. To this end, we computed the mean of gene expression values belonging to different individuals for each gene and organ. In order to compute expression divergence between each pair of species, we then calculated the difference between the mean expression values of the orthologous gene pairs, normalized by the sum of the mean expression values in a given organ. We then partitioned these pairwise expression differences into two subsets according to whether orthologous genes did or did not contain TRs in their promoters. We observed a significant increase in pairwise expression differences when genes have TRs in their promoters. More specifically, between human and chimpanzee orthologs with repeats within 5kb upstream of their TSS showed higher mean expression difference (0.264) compared to those without repeats (0.257, P < 10^-6^, based on Wilcoxon Rank Sum Test (Mann and Whitney 1947). Similarly, human-macaque orthologs (P < 0.01, for all organs) and chimpanzee-macaque orthologs (P < 10^-5^, for all organs) with TRs showed a higher mean expression difference (0.250 and 0.254, respectively) than orthologous genes without repeats (0.243 and 0.242, respectively).

In order to avoid noise and bias for organ-specific gene expression variation differences, we next took a phylogenetic approach and performed a bootstrap-like resampling analysis, where gene expression values were sampled from different individuals of a species (see Methods). We computed two different expression distance matrices of (1000 replicates) × (3 species pairs) for each organ and employed these matrices to construct neighbor-joining gene expression trees. Except for the macaque branch for liver- and heart-specific expression trees, all branches were significantly longer for repeat-containing genes in both species (P < 10^-10^; based on a t-test with n = 1000, df = n-1, throughout unless otherwise mentioned) (Supplementary Figure S3). The total tree length of genes with repeats was significantly greater in all organs (P < 10^-200^ except for liver, where P = 0.02) (Figure 4A). These observations also held when we changed the length of the upstream regions we considered. Repeat-containing genes diverged more rapidly in their expression, and this difference was most pronounced for repeats within 1kbp upstream of the transcription start site (95% CI: 0.0145, 0.0146). The difference got progressively smaller as we included repeats that are further away from the transcription start site (95% CI for windows of length 10 kbp: 0.003, 0.006; 15 kbp: 0.0021, 0.0024; 20 kbp: 0.0004, 0.0007, Supplementary Figure S4).

**Figure 4.**
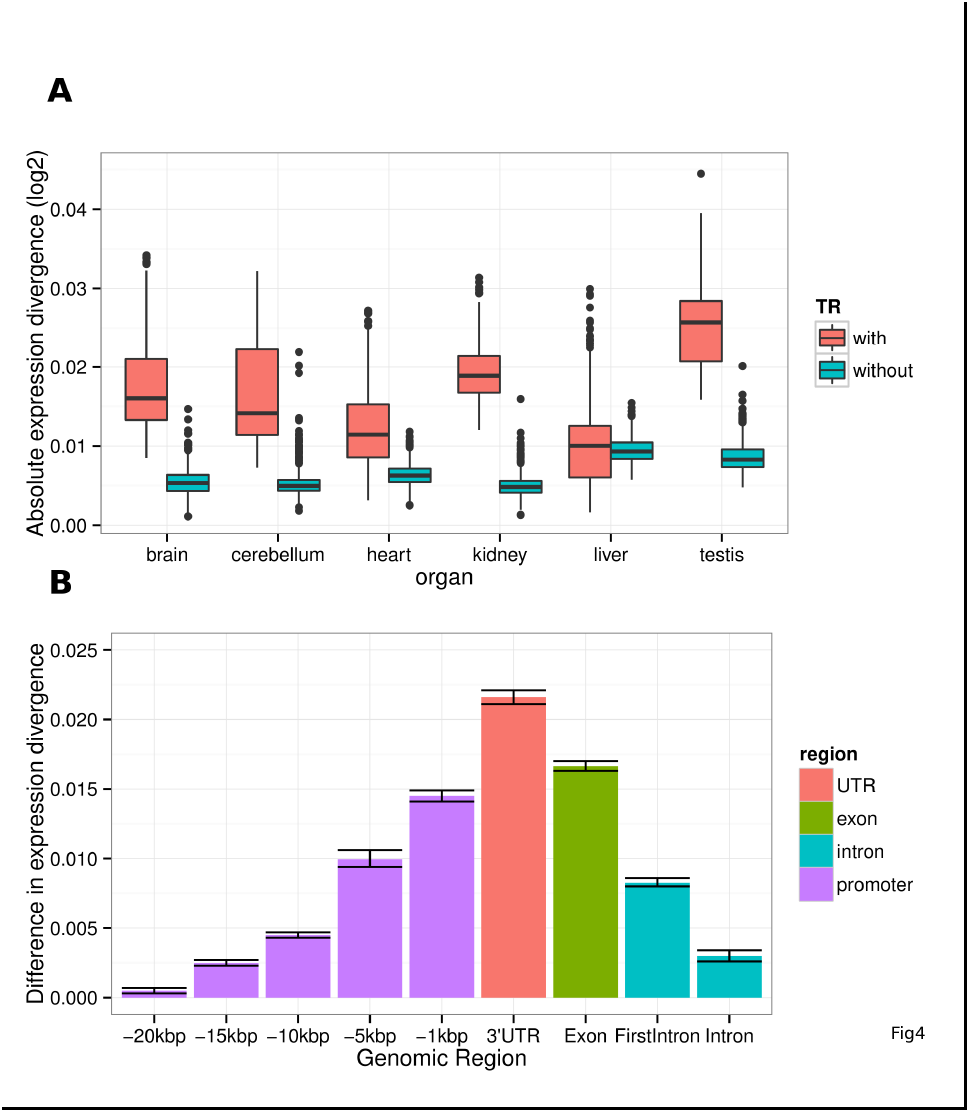
Relationship between expression divergence and the presence of repeats in gene promoters and other genic regions. A) Boxplot of total tree lengths of genes with repeats (thick lines) and genes without repeats (thin lines). Horizontal lines in the middle of each box mark the median, edges of boxes correspond to the 25th and 75th percentiles, and whiskers cover 99.3% of the data points. B) Presence of tandem repeats associate with higher expression divergence. Bars present mean differences in expression divergence, based on pairwise expression tree length differences between repeat-containing and non-repeat-containing genes. Repeats found in upstream regions of length 20kbp, 15kbp, 10kbp, 5kbp, 1 kbp, as well as in 3’UTRs, exons, first introns and all introns were considered, as indicated on the horizontal axis. Note that all expression differences are positive, indicating that repeat-containing genes, regardless of category, diverged more rapidly. Whiskers represent 95% confidence intervals.

Since gene expression levels may play a role in expression divergence (Lehner 2008; Macneil and Walhout 2011; Pilpel 2011) we then asked whether the association between TRs and expression divergence varies with expression level. To this end, we distinguished between highly and lowly expressed genes, choosing the median expression level of all genes among all individuals in a species (and separately for each organ) as a threshold for high and low expression. We followed our previous randomization procedure and found that TRs associate with expression divergence to a similar extent for highly expressed genes, (P < 10^-117^, all organs considered together; n = 6000; df = n-1) as they do for lowly expressed genes (P < 10^-129^, N = 6000). CpG islands play an important role in mammalian gene regulation (Saxonov et al. 2006) and may thus affect gene expression divergence. Therefore it is important to note that these observations also held when we controlled for repeats overlapping with CpG islands. More specifically, we found that only 215 of all identified tandem repeats overlapped with a CpG island (see Supplementary Text S1 for details). When analyzing expression divergence while excluding those overlaps, we found that the presence of repeats was still strongly associated with expression divergence (P < 10^-178^, for all organs except for liver, where P < 10^-6^, n = 6000).

Repeats in other potential regulatory regions also have an influence on the divergence of gene expression. Specifically, genes containing TRs within 1 kb of the 3’ UTR also showed significantly greater expression divergence in all organs except the testis (P < 10^-47^, for all organs; Supplementary Figure S5A). Moreover, those 2,468 human genes with exon-containing repeats also showed greater expression divergence in all organs (P < 10^-269^, for all organs; supplementary figure S5B). Finally, repeats in introns were also associated with greater expression divergence (P < 10^-198^) for all organs except for the heart (P <10^-85^) and the liver (P < 10^-292^), both of which show opposite patterns (Supplementary figure S5D). The mean difference of tree lengths for repeats found in any intron was smaller compared to the mean difference for the repeats found in first introns (95% CI: 0.0080, 0.0086). Overall, our gene expression trees indicate greater expression divergence between genes without repeats and genes that contain repeats, in the following order of decreasing divergence: repeats in 3’ UTR regions (mean difference in tree lengths: 0.022) >> repeats in exons (0.017) >> repeats in promoters (0.015) >> repeats in 1st intron (0.008) >> repeats in any intron (0.003) (Figure 4B).

To check to which extent polymorphism within species might be affecting our results, we used our set of genotyped TRs, and classified repeats either as polymorphic or fixed (no variation in copy number of repeat unit) within each taxon. To this end, we used our genotyped TR data in the panel of 27 humans and 16 chimpanzees. We found that for all tissues, genes with promoters containing repeats observed to be polymorphic in both taxa, showed significantly more divergence than (i) genes with the same repeat genotype (repeat length) in both taxa and (ii) genes without repeats (P < 10^-168^ and P < 10^-163^, respectively, based on Wilcoxon Rank Sum Test) (Figure 5).

**Figure 5.**
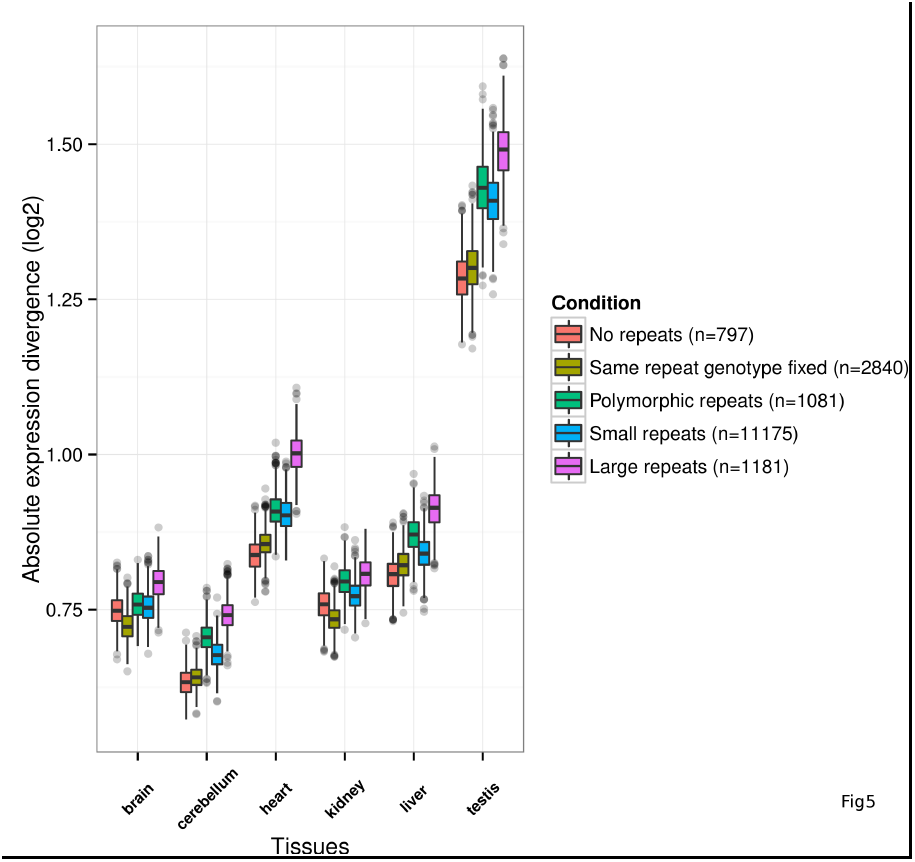
Relationship between expression divergence and within-species repeat genotype conservation in human and chimpanzees. Boxplots produced by resampling 1000 data points each corresponding to the average log2-transformed expression divergence value between human and chimpanzee for a particular tissue and for genes associated to a particular category. The tissues are shown on the x axis, and the y axis corresponds to the absolute mean expression divergence between humans and chimpanzees. The categories considered are: genes with no repeats in promoters (red); genes containing exclusively repeats in promoters which have the same repeat length fixed across all human and chimpanzee samples (yellow); genes containing exclusively repeats which are polymorphic in human and chimpanzees (green); genes containing small repeats (repeat unit length of 1-5 bp and less than 100bp total repeat length) in their promoter (blue), and genes containing large repeats (repeat unit length of 2-50 bp) in the promoter. Genes lacking repeats in promoters or repeats for which the same repeat genotype length is found across human and chimpanzee samples show the least amount of expression divergence for all tissues.

Using the genotyping data for human and chimpanzee, as well as for the other three primate taxa (bonobos, gorillas and orangutans), we finally checked for overrepresented biological process terms in genes that contained repeats in three categories: (i) genes whose repeat genotype length was different between human and chimpanzees and fixed within each taxon (n=2804), (ii) genes whose repeats were the same and fixed in all genotyped nonhuman primates but polymorphic in humans (n=1754), and (iii) genes whose repeats are fixed in humans but polymorphic in all genotyped nonhuman primates (n=2178). We found that for the first two categories processes related to cell-adhesion, neurogenesis and neural development are enriched, while for category (iii) processes related to detection and response to chemical and biotic stimuli, sensory perception of taste and smell, and skin development are enriched (Supplementary Table S5).

## DISCUSSION

TRs are abundant in human and nonhuman primate genomes, but their variation in natural populations and their potential regulatory role remain largely unexplored. Recent advances in sequencing technology, together with the development of computational tools, have finally made it affordable to accurately genotype up to millions of TRs at the genome-wide level, and to circumvent much of the constraints that traditional repeat genotyping methods entail (Gymrek et al. 2012; Highnam et al. 2013; Guilmatre et al. 2013; Duitama et al. 2014; Willems et al. 2014; Carlson et al. 2014).

Here, we surveyed the genome-wide diversity of TRs in a large collection of high quality human and nonhuman great ape genomes, and showed for the first time their impact on gene expression divergence between human and other primates. We observed a strong association between polymorphic repeats in gene promoters and increased expression divergence, an observation that was robust to changes in the method used to identify tandem repeats and to assess gene expression divergence. This association existed for most of all organ-specific expression data, except in some cases for testis and liver. Distinct expression patterns in these organs have also been observed by others (Hsieh et al. 2003; Somel et al. 2008; Brawand et al. 2011) in different contexts.

Repeats closer to the transcription start site were associated with greater expression divergence, an observation that might be explained through core promoter modules occurring preferentially close to this site and exerting a strong influence over transcriptional regulation (Wray et al. 2003; Spitz and Furlong 2012). Moreover, an association with expression divergence held also for repeats in other genic regions. The strongest of them was evident for 3’UTRs, consistent with their known role in gene regulation (Yoon et al. 2012). In addition, repeats in first introns were associated with greater expression divergence than repeats in other introns. This observation is consistent with previous work showing that most intronic regulatory regions occur in the first intron (Rohrer and Conley 1998), that the first intron has the highest divergence between human and chimps (Gazave et al. 2007), and that the first intron influences gene expression more than others (Jonsson et al. 1992; Charron et al. 2007).

Our analysis of expression divergence in repeats that could be genotyped at the population level also supported a relationship between differential repeat length in promoters and higher gene expression divergence in humans and chimpanzees. Our results indicated that polymorphic repeats may have a significant impact on the levels of expression divergence, particularly when compared to genes without repeats, or genes in which the same repeat genotype is fixed across human and chimpanzee samples, thus suggesting that repeat variation may elicit changes in gene expression levels across species. Intriguingly, genes containing repeats whose genotype is conserved across nonhuman primates but polymorphic or fixed for a different genotype in humans seem to be enriched for functions related to neurogenesis and neural diffentiation, as well as development of the nervous system and cell-adhesion. Furthermore, genes for which all repeats are fixed in humans, but not fixed in other nonhuman primates seem to be enriched for processes related to stimulus detection, sensory perception, and skin development. Such biological processes may be associated with the evolution of human specific cognitive traits and the response to new environments, respectively. However, since the nature of these fixation events is not certain, it will be important in the future to distinguish between the demographic (e.g., human population bottlenecks) and selective contributions in order to realize the true significance of these findings. Moreover, these results provide further motivation for future studies to clarify the exact role of these genes in primate evolution, and the extent to which repeats may have been involved in their regulation.

The importance of doing so is also in part supported by the findings of previous studies that explored and found a clear relationship between repeat variation and phenotypical or gene expression divergence. Accumulating evidence from exhaustive genetic studies have shown that TR variation has dramatic, often background-dependent phenotypic effects in model organisms (Verstrepen et al. 2005; Kashi and King 2006; Fondon et al. 2008; Borel et al. 2012; Raveh-Sadka et al. 2012; Morrison et al. 2012; Egbert and Klavins 2012). In yeasts, TR variation in promoters has been shown to alter gene expression (Vinces et al. 2009). An especially remarkable example in mammals, regards features of a dog’s snout, such as the degree of dorsoventral nose bend and midface length, which correlate with the ratio of the length of two tandem repeats in a gene that regulates bone formation (Fondon and Garner 2004). Repeat polymorphisms have also been linked to behavioural and cognitive functions. For example, the presence of tandem repeats in the 5’ untranslated region of the vole vasopressin 1a receptor gene correlates with social behavior (Fondon et al. 2008). Mutations in this gene have been implicated in autism (Kim et al. 2002). The gene’s orthologue in chimpanzees has a partial deletion in those tandem repeats, whereas the more social human and bonobo (Waal 2009) have the complete set of repeats (Hammock and Young 2005). Furthermore, a recent study (Hellen and Kern 2015), has showed an enrichment for insertions in human genes and gene regulatory regions associated with recent human evolution. More specifically, they found that TR insertions are significantly more frequent thatn expected among other type of insertions, and that they are mostly fixed in the human lineage. Taken together with all other studies, our observations further suggest an important contribution of TRs in primate gene expression evolution.

In this manuscript, we also showed how genome wide short TRs genotyped from whole genome sequencing data provide a valid means to uncover substructure and divergence patterns in human populations and great ape species, by showing that they agreed with previous surveys in human and nonhuman great apes based on single-nucleotide polymorphisms (Li et al. 2008; Prado-Martinez et al. 2013). These results have several implications for conservation and breeding programs of great apes, since they provide several new loci whose genotypes can be used to distinguish among different taxonomical groups or populations, and to assess the degree of diversity present in natural populations (McMahon et al. 2014). Importantly, due to their high polymorphism, TRs present several advantages relative to other popular molecular markers. In particular, their higher mutation rate means that a low number of markers can be used, an advantage for conservation studies, which often use highly degraded non-invasive samples.

Several limitations are associated with this work. First, repeat content, repeat length, and repeat unit sizes may be affected by different mutation rates. While one would ideally want to take these differences into account, doing so would have limited our statistical power to detect structure patterns at several levels. Another limitation stems from whole genome sequencing genotyping with short read technologies and regards the maximum length of a repeat that can be genotyped. The reason is that genotyping a repeat requires it to be wholly encompassed by any given short read. Because we wanted to analyze only non-overlapping repeats, the sequence reads of ∼100 base pairs used in this study imply that we were limited to the analysis of only ∼58% of the total fraction of TRs we identified on the human reference genome using Tandem Repeat Finder. Nonetheless, we were still able to genotype thousands of TRs, and assess their conservation within and across different primate natural populations.

In the future, studies of the repeat landscape will be facilitated not only by the use of longer reads, but also by a thorough subsequent genotyping of a subset of repeats, using one of the recently developed methods that specifically target repeats (Guilmatre et al. 2013; Duitama et al. 2014; Carlson et al. 2014). These methods rely on a pre-sequencing enrichment step for repeats. They are currently able to target and genotype several thousand repeats in many individuals, and do so in a much more accurate fashion than *in silico* methods. The combination of the two approaches will yield much reliable and important information regarding repeat variation in natural populations. Such TR genotyping from whole genome sequencing data will have a profound impact on many fields, from conservation genetics to forensics, and in elucidating the role of TRs in complex trait heritability (Press et al. 2014).

In a seminal paper, King and Wilson (King and Wilson 1975) observed about humans and chimpanzees that “their macromolecules are so alike that regulatory mutations may account for their biological differences.” Since then, we have learned that such mutations, and in particular mutations that cause gene expression change, are indeed important in the evolution of primates and other organisms (Wren et al. 2000; Stranger et al. 2005, 2007; Fondon et al. 2008; Dimas et al. 2009; Vinces et al. 2009; Gemayel et al. 2010). Our work shows that TRs, a type of sequence with unusually high mutability, may be an important class of regulatory mutations that are responsible for such species differences.

## DATA ACCESS

Raw sequence data from published genomes are available through the Sequence Read Archive (SRA) (SRP018689, SRP009145, SRP001139, SRP001703), and at https://www.simonsfoundation.org/life-sciences/simons-genome-diversity-project-dataset/. The 9 newly sequenced African genomes have been deposited at the SRA database (SRP052818).

## METHODS

### Genomic sequence data

We used a total of 83 samples sequenced with Illumina paired-end reads at high-coverage (>20X) and with read length ranging from 50–102-base pairs (bps). Specifically, we used both publicly available datasets and newly sequenced human genomes. For the non-human primates, we used the complete collection from the Great Apes Genomic Project (Prado-Martinez et al. 2013). In addition, we selected a total of 27 human male samples, 14 from the Human Genome Diversity Project dataset and 2 Dinka individuals (Meyer et al. 2012; Prüfer et al. 2014), 9 African genomes (Lorente et. al 2015 in preparation), and two complete genomes from a Yoruba individual (Li et al. 2011) and a Khoisan individual (Schuster et al. 2010) (See Supplementary Table S6 for sample information). All reference genomes for human (hg19), chimpanzee (panTro2.14) (Sequencing and Consortium 2005), gorilla (gorGor3) (Scally et al. 2012), and orangutan (ponAbe2) (Locke et al. 2011) were retrieved from the UCSC genome browser (Kent et al. 2002).

### Tandem Repeat Identification

For the genomic diversity data analysis we used the set of tandem repeats that come with the Repeatseq (Highnam et al. 2013) software. This is a set containing tandem repeats with a repeat unit between 1 and 5 base pairs in length, identified in the human reference genome (hg19) using Tandem Repeat Finder (TRF) (Benson 1999) v2.30, with parameters “2 5 5 80 10 14 5”, and further filtered so that any two repeats are at least 21 base pairs apart.

These parameters identified all repeats with TRF score ≥14, i.e. repeats with a total length of at least 7 bps when the repeat unit is a single base pair and 6 bps for all other repeat unit sizes considered (2 to 5 bps), and which passed some statistical criteria based on the total length of the sequence being analyzed, the sequence match and indel probabilities relative to the repeat unit, and the repeat unit size (see Supplementary Table 7 for more information on the parameter values and their description).

Using the same parameters, we also identified tandem repeats in the panTro2.1.4, gorGor3 and ponAbe2 reference genomes, corresponding respectively to chimp, gorilla and orangutan.

Additionally, we also identified tandem repeats occurring only in or near genes, using more stringent parameters and allowing for larger repeat motifs. These included the promoter (5,000 base pairs (bps) upstream from the transcription start site, unless stated otherwise), exons, the first intron, all introns, and the 3’ untranslated region (1000 bps downstream from each gene’s stop codon). Here, we considered both micro- and minisatellites with tandem repeat units up to 50 nucleotides in length. Longer repeats are less variable and therefore less likely to cause phenotypic divergence (Li et al. 2002, 2004; Kelkar et al. 2008; O’Dushlaine and Shields 2008; Payseur et al. 2011). Specifically, for this dataset we only considered repeats with TRF scores ≥80, i.e. repeats with a total length of at least 40 nucleotides (e.g., 10 copies of a tetranucleotide repeat or 20 copies of a dinucleotide repeat), an incidence of indels in adjacent repeat units ≤10% (e.g., a di nucleotide repeat with ten copies can have up to two indels overall relative to the adjacent pattern), and sequence identity of adjacent repeat units at least 90% (e.g., at least 18 nucleotides of a di nucleotide repeat of ten copies must match the adjacent pattern). One motivation for these stringent thresholds is that the variability of tandem repeats increases strongly for repeats of high sequence similarity and TRF Scores (O’Dushlaine and Shields 2008).

### Identification of species-specific repeats at reference genome level

We first sought to retrieve the sequences corresponding to the set of tandem repeat loci identified respectively in the human, chimpanzee, gorilla, and orangutan reference genomes with Tandem Repeat Finder (TRF, See *Tandem Repeat Identification*), in each of the other three reference genomes. To do this, we first downloaded the Enredo, Pecan, Ortheus (EPO) alignment (Paten et al. 2008) (downloaded from ftp://ftp.ensembl.org/pub/current_emf/ensembl-compara/epo_6_primate/), a multiple sequence alignment computed at the reference genome level between six primate species (including the four used here), and then used a python script (available at https://bitbucket.org/james_taylor/bx-python/) to retrieve the sequences corresponding to each repeat locus that had been identified in a particular reference genome. We used this multi-alignment information in order to identify and then remove repeat loci for which in any of the taxa, the corresponding sequence portion was missing from their reference genome, was duplicated, or was missing sequence information (nucleotides marked as “N”). Next, we ran TRF on the retrieved sequences, and considered them to contain a repeat if either the repeat motif identified in the original tandem repeat list was the same as the one identified now by TRF, or if this particular repeat motif was found at least twice in tandem, and the total repeat length was at least four base pairs long. We then looked for repeat loci whose coordinates overlapped on the same original reference genome where these repeats had first been identified, and removed in each case all but one locus, that for which repeats were identified across the largest number of reference genomes (taxa). Lastly, we computed the number of repeat loci in each reference genome according to whether the corresponding sequences in the other reference genomes contained a repeat or not. We performed this entire procedure for all four reference genomes.

### Mapping and genotyping of tandem repeats

For each sample we first mapped 50,000 reads to the human reference genome (hg19) using Novoalign aligner version 2.08.02 (http://www.novocraft.com), in order to calculate the mean and standard deviation of the insert size between the reads in each set of paired-end reads. We then supplied this information to the aligner when performing the mapping of all reads to the human genome. Upon mapping we realigned reads located around indels using the Genome Analysis Toolkit (GATK) version 2.5 (McKenna et al. 2010), and used Picard Tools v1.7 (http://picard.sourceforge.net/) for removal of duplicate reads. Finally, using the set of tandem repeats identified with Tandem Repeat Finder (Benson 1999) we genotyped all samples using Repeatseq version 0.8.2 (Highnam et al. 2013), with default parameters and setting the “-emitconfidentsites” option.

### Validation of tandem repeats identified in nonhuman primates

From each of the chimpanzee, gorilla, and orangutan sample sets, we selected one individual sample. We then mapped each of these samples to its own species reference (panTro 2.1.4, gorGor3 and ponAbe2 respectively for chimpanzee, gorilla and orangutan) using the Novoalign aligner v2.08.02 (http://www.novocraft.com), and after duplicate removal with Picard (http://picard.sourceforge.net/), genotyped repeats with Repeatseq (Highnam et al. 2013) on the set of tandem repeats identified by running Tandem Repeat Finder (Benson 1999) in the sample's species reference genome. We then used the UCSC liftover tool (Hinrichs et al. 2006) to get the positions of the genotyped repeats in the hg19 human reference genome, and checked if the genotypes called with this particular approach and the standard one (using the hg19 human reference) agreed. For each species, positions where the genotypes were found to disagree were excluded from further analysis. Due to the high degree of similarity between bonobos and chimpanzees, the set of positions excluded for chimpanzee are the same that were excluded for bonobo.

### Genotype data filtering and tandem repeat copy-number definition

For each sample we kept only those genotyped tandem repeats covered by at least 10 reads. In addition, we removed any genotypes which in the genomic context of a particular species are theoretically impossible to occur biologically, i.e. heterozygous genotypes in the X and Y sex chromosomes of male individuals, or any genotype called in the Y sex chromosome for female individuals. Furthermore, except for the analyses of heterozygosity and repeat copy number difference relative to human reference genome, we only considered tandem repeats where dividing the repeat length by the repeat unit length resulted in an integer.

In all analyses genotypes are represented by the absolute copy number of the repeat unit in each allele, calculated as the tandem repeat total length divided by the repeat unit length.

### Absolute repeat copy-number distance from human reference genome

We estimated absolute repeat copy number distance from the human reference genome for groups corresponding to all chimpanzee and gorilla subspecies, bonobos, the two orangutan species, and for three groups of humans, namely African hunter-gatherers, all Africans that did not fit in the former category, and non-Africans. For all repeat loci genotyped, we first subtracted the repeat copy number found by TRF in the human reference genome from the repeat copy number of each allele. We then resampled both TR loci and samples of each group one hundred times. More specifically, in each resampling step, 500,000 TR loci were randomly sampled for each taxon (bonobo and chimpanzees were considered as a single taxon), for which three individuals, also randomly sampled, had all been genotyped. For each of these repeat loci and sets of three samples, we averaged the absolute values of ten randomly sampled alleles corresponding to the repeat copy number differences relative to the human reference genome. We then added 0.15 to each one of these averages and kept only the integer part of the resulting number (e.g. if at a particular locus the average repeat copy number difference relative to the human reference of the 10 alleles sampled across Africans was 2.87, we would consider their repeat copy number difference relative to the human reference to be 3, since the integer part of 3.2 (2.87+0.15=3.2) is 3 in that case). We quantified the number of times each repeat copy number difference was observed within each group in each of the 100 iterations, and plotted these results with the geom_smooth function of ggplot (Wickham 2009), which performs a local regression fit to the data.

### Heterozygosity

To estimate heterozygosity, we used the set of one hundred re-samplings of 500,000 repeat loci and three samples per group, which had been generated previously in the analysis concerning the absolute repeat copy-number distance from the hg19 reference. Then, for each group and locus, we randomly sampled fifty pairs of alleles from a pool consisting of up to three samples, classifying each pair either as being homozygous or heterozygous, if either the two alleles were the same or different, respectively. Finally we divided the total number of heterozygotes found, by the total number of loci analyzed, and calculated the average of this estimate for each of the 100 re-samplings. We then used the 100 averages obtained per group to produce the boxplots for each group.

### Population structure analysis

For each species we selected a set of positions where all individuals had been genotyped, and for which all genotyped positions are at least 2000 base pairs apart. Then, for each set we proceeded to perform a principal component analysis (PCA), after mean centering the data, using the adegenet R package (Jombart 2008; Jombart and Ahmed 2011).

Using the same set of positions, we also used STRUCTURE (Pritchard et al. 2000) version 2.3.4 to look at the genetic structure present within each species. For this we selected an admixture model with correlated allele frequency, and for each K going from 2 to 7, where K is the number of putative ancestral clusters characterized by a subset of allele frequencies. All runs were performed with 1,000,000 Markov Chain Monte Carlo repeats, following a burn-in period of 10,000 iterations. W For each species and K we then repeated the same analysis two additional times, to ensure consistency of the results, and examined a time series of the alpha values, in order to make sure that convergence on alpha was reached following the burn-in period.

In order to find, for each species, the number K of genetically meaningful clusters that might exist within each species, we calculated Evanno's delta K (Evanno et al. 2005), using a stand-alone script (Earl and VonHoldt 2011).

### Genetic differentiation

The pairwise distance between all samples, defined by Rst, an analogue to Wright's Fst, was calculated both for individuals of each species separately using microsat (http://genetics.stanford.edu/hpgl/projects/microsat/).

### Ancestrally informative markers

For each species of nonhuman primates we first selected those loci for which at least 75% of the individuals were genotyped both at the subspecies and species level in chimpanzees and gorillas, and at the species level in orangutans. We then identified those loci which contain both a repeat genotype and alleles unique to each chimpanzee and gorilla subspecies, as well as orangutan species.

### Tandem repeats genomic context

We characterized TRs according to whether they were located within promoters, splice sites, 3 or 5 UTR's, exons or introns, using the GenomicRanges package version 1.14.4 (Lawrence et al. 2013).

### Repeat genotype conservation

We classified TRs in each taxon as being fixed or polymorphic across the genotyped samples. For a TR to be considered as fixed within a taxon, we required that over 40% of the samples had been genotyped, and that at least 95% of these share the same repeat genotype length. For a TR to be considered polymorphic, it needed to have at least four samples genotyped, with at least two different genotypes present. We classified each repeat locus as polymorphic or fixed in both human and nonhuman primates.

### Gene ontology

We used the topGO R package (Alexa and Rahnenfuhrer 2010) to search for gene ontology terms related to biological processes which may be enriched both in human specific repeats identified at the reference level, and for particular TR genotype conservation categories within taxa (fixed or polymorphic). We used a Fisher's test to infer this overrepresentation, followed by an adjustment for multiple testing of the p-values produced using the Benjamini-Hochberg method (Kasen et al. 1990).

### Gene expression and sequence data

The gene expression data we used were based on RNA sequencing of ∼3.2 billion 76-base pair Illumina Genome Analyser IIx reads (Brawand et al. 2011). Expression levels are indicated as log2-transformed reads per kilobase pair of exon model per million mapped reads. This provided one-to-one gene expression measurements from multiple primates, where each gene's expression had been measured in six different organs (brain, cerebellum, heart, kidney, liver, testis) for between 1 and 6 individuals per species. From this data set, we used RNA-seq based expression values of all 13,035 one-to-one gene orthologs from humans, chimpanzees and macaques. We obtained DNA sequences of the genes in our expression data set through the Biomart tool of Ensemble (Kinsella et al. 2011), using human annotation version GRCh37.p10, chimpanzee annotation CHIMP2.1.4, and macaque annotation MMUL_1.

### Gene expression divergence

The gene expression data set we used (Brawand et al. 2011) contained gene expression measurements from several individuals of a species for each gene and organ. We took advantage of this fact to assess statistical differences in gene expression divergence with a bootstrap-like resampling procedure, where we sampled gene expression values from different individuals of a species to create 1000 replicate data sets (n=13,035) for each organ, and species.

We partitioned gene pairs in each such data set into two groups: gene pairs where genes of a given species contained tandem repeats in a specific region of interest, such as a promoter, and gene pairs without such repeats. We then computed, separately for genes in the two groups, a pairwise matrix of Euclidean gene expression distance between all genes in a pair of species, based on the formula (Tirosh et al. 2006):

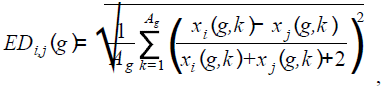
where i and j stand for species i and j (e.g. human and chimpanzee), g is a (binary) indicator variable reflecting which of two sets of genes (with or without repeats) are analyzed, Ag is the number of genes in that set, k is a gene-specific index and x is the expression level of a gene. To give an example, *x_human_*^(*norepeat*, 1)^ is the gene expression value of the first gene in the human gene set without repeats for a given organ, and ED_human, macaque_ (repeat) is the expression distance between repeat-containing human and macaque gene pairs for a given replicate data set.

Overall, we created 12 separate expression distance matrices of size (1000×3), for two gene subsets based on repeat presence and for six organs. We used these matrices to construct gene expression trees using the neighbor-joining approach (implemented in the ‘ape’ package (Paradis et al. 2004) in R (http://www.R-project.org/)). We used the branch lengths of the trees we constructed as a measure of gene expression divergence. To test the null-hypothesis that the expression divergences (branch lengths) of the 1000 sampled trees were significantly different between the two gene subsets for each organ, we used paired t-tests (N=1000, df=n-1 unless otherwise mentioned). All P values are reported after Bonferroni correction (Dunn 1961) for multiple testing and they were robust to number of bootstrap replicates. We performed all statistical analyses using MATLAB (7.10.0, The MathWorks Inc., Natick, MA, R2010a).

### Expression divergence versus repeat genotype conservation

We used again the expression data available for genes present both in human and chimpanzee (Brawand et al. 2011), and calculated the mean absolute expression divergence between human and chimp for each tissue and TR genotype conservation category by randomly sampling 1000 genes with replacement. For each of these genes we sampled the expression for both species from among the samples (up to 6 by tissue) with expression different from zero, and calculated the log2 of their absolute expression levels. We then ranked these by their absolute expression divergence level, and averaged their value after removing the top and bottom 25% values. This step was repeated 100 times, and the values used to generate the boxplot.

To test for significance across different categories comparisons, we used the wilcox.test R function to implement a Wilcoxon-Mann-Whitney statistical test.

## ACKNOWLEDGMENTS

AW and TBS acknowledge support through Swiss National Science Foundation grant 31003A_146137, as well as through the University Priority Research Program in Evolutionary Biology at the University of Zurich. TBS thanks Maria Anisimova for helpful discussions. TC thanks Irene Hernando-Herraez, Javier Prado-Martinez, and Rui Faria for their help and constructive comments. TMB was supported by an ERC Starting Grant (260372) and MICINN (BFU2011-28549, Spain).

## REFERENCES

Alexa A, Rahnenfuhrer J. 2010. topGO: topGO: Enrichment analysis for Gene Ontology. R package version 2.18.0. *October*.

Arora N, Nater A, van Schaik CP, Willems EP, van Noordwijk MA, Goossens B, Morf N, Bastian M, Knott C, Morrogh-Bernard H, et al. 2010. Effects of Pleistocene glaciations and rivers on the population structure of Bornean orangutans (Pongo pygmaeus). Proc Natl Acad Sci U S A 107: 21376–21381.

Becquet C, Patterson N, Stone AC, Przeworski M, Reich D. 2007. Genetic structure of chimpanzee populations. PLoS Genet 3: e66.

Benson G. 1999. Tandem repeats finder: a program to analyze DNA sequences. Nucleic Acids Res 27: 573–580.

Bergl RA, Vigilant L. 2007. Genetic analysis reveals population structure and recent migration within the highly fragmented range of the Cross River gorilla (Gorilla gorilla diehli). Mol Ecol 16: 501–16.

Borel C, Migliavacca E, Letourneau A, Gagnebin M, Béna F, Sailani MR, Dermitzakis ET, Sharp AJ, Antonarakis SE. 2012. Tandem repeat sequence variation as causative cis-eQTLs for protein-coding gene expression variation: the case of CSTB. Hum Mutat 33: 1302–9.

Bowden R, MacFie TS, Myers S, Hellenthal G, Nerrienet E, Bontrop RE, Freeman C, Donnelly P, Mundy NI. 2012. Genomic tools for evolution and conservation in the chimpanzee: Pan troglodytes ellioti is a genetically distinct population. PLoS Genet 8: e1002504.

Brawand D, Soumillon M, Necsulea A, Julien P, Csárdi G, Harrigan P, Weier M, Liechti A, Aximu-Petri A, Kircher M, et al. 2011. The evolution of gene expression levels in mammalian organs. Nature 478: 343–8.

Brinkmann B, Klintschar M, Neuhuber F, Hühne J, Rolf B. 1998. Mutation rate in human microsatellites: influence of the structure and length of the tandem repeat. Am J Hum Genet 62: 1408–15.

Carlson KD, Sudmant PH, Press MO, Eichler EE, Shendure J, Queitsch C. 2014. MIPSTR: a method for multiplex genotyping of germ-line and somatic STR variation across many individuals.

Charron M, Chern J-Y, Wright WW. 2007. The cathepsin L first intron stimulates gene expression in rat sertoli cells. Biol Reprod 76: 813–24.

Choi JK, Kim Y-J. 2008. Epigenetic regulation and the variability of gene expression. Nat Genet 40: 141–7.

Dimas AS, Deutsch S, Stranger BE, Montgomery SB, Borel C, Attar-Cohen H, Ingle C, Beazley C, Gutierrez Arcelus M, Sekowska M, et al. 2009. Common regulatory variation impacts gene expression in a cell type-dependent manner. Science 325: 1246–50.

Duitama J, Zablotskaya A, Gemayel R, Jansen A, Belet S, Vermeesch JR, Verstrepen KJ, Froyen G. 2014. Large-scale analysis of tandem repeat variability in the human genome. Nucleic Acids Res 42: 5728–41.

Dunn OJ. 1961. Multiple Comparisons Among Means. J Am Stat Assoc 56: 52.

Earl D a., VonHoldt BM. 2011. STRUCTURE HARVESTER: a website and program for visualizing STRUCTURE output and implementing the Evanno method. Conserv Genet Resour 4: 359–361.

Egbert RG, Klavins E. 2012. Fine-tuning gene networks using simple sequence repeats. Proc Natl Acad Sci U S A 109: 16817–22.

Ellegren H. 2004. Microsatellites: simple sequences with complex evolution. Nat Rev Genet 5: 435–45.

Enard W, Paabo S. 2004. Comparative primate genomics. Annu Rev Genomics Hum Genet 5: 351–378.

Evanno G, Regnaut S, Goudet J. 2005. Detecting the number of clusters of individuals using the software STRUCTURE: a simulation study. Mol Ecol 14: 2611–20.

Fondon JW, Garner HR. 2004. Molecular origins of rapid and continuous morphological evolution. Proc Natl Acad Sci U S A 101: 18058–63.

Fondon JW, Hammock E a D, Hannan AJ, King DG. 2008. Simple sequence repeats: genetic modulators of brain function and behavior. Trends Neurosci 31: 328–34.

Fünfstück T, Arandjelovic M, Morgan DB, Sanz C, Breuer T, Stokes EJ, Reed P, Olson SH, Cameron K, Ondzie A, et al. 2014. The genetic population structure of wild western lowland gorillas (Gorilla gorilla gorilla) living in continuous rain forest. Am J Primatol 76: 868–78.

Gazave E, Marqués-Bonet T, Fernando O, Charlesworth B, Navarro A. 2007. Patterns and rates of intron divergence between humans and chimpanzees. Genome Biol 8: R21.

Gemayel R, Vinces MD, Legendre M, Verstrepen KJ. 2010. Variable tandem repeats accelerate evolution of coding and regulatory sequences. Annu Rev Genet 44: 445–77.

Gonder MK, Locatelli S, Ghobrial L, Mitchell MW, Kujawski JT, Lankester FJ, Stewart C-B, Tishkoff S a. 2011. Evidence from Cameroon reveals differences in the genetic structure and histories of chimpanzee populations. Proc Natl Acad Sci U S A 108: 4766–71.

Goossens B, Chikhi L, Ancrenaz M, Lackman-Ancrenaz I, Andau P, Bruford MW. 2006. Genetic signature of anthropogenic population collapse in orang-utans. PLoS Biol 4: e25.

Greminger MP, Stölting KN, Nater A, Goossens B, Arora N, Bruggmann R, Patrignani A, Nussberger B, Sharma R, Kraus RHS, et al. 2014. Generation of SNP datasets for orangutan population genomics using improved reduced-representation sequencing and direct comparisons of SNP calling algorithms. BMC Genomics 15: 16.

Gross DS, Garrard WT. 1988. Nuclease hypersensitive sites in chromatin. Annu Rev Biochem 57: 159–197.

Guilmatre A, Highnam G, Borel C, Mittelman D, Sharp AJ. 2013. Rapid multiplexed genotyping of simple tandem repeats using capture and high-throughput sequencing. Hum Mutat 34: 1304–11.

Gymrek M, Golan D, Rosset S, Erlich Y. 2012. lobSTR: A short tandem repeat profiler for personal genomes. Genome Res 22: 1154–62.

Hammock E a D, Young LJ. 2005. Microsatellite instability generates diversity in brain and sociobehavioral traits. Science 308: 1630–4.

Hellen E, Kern A. 2015. The role of DNA insertions in phenotypic differentiation between humans and other primates. Genome Biol Evol.

Highnam G, Franck C, Martin A, Stephens C, Puthige A, Mittelman D. 2013. Accurate human microsatellite genotypes from high-throughput resequencing data using informed error profiles. Nucleic Acids Res 41: e32.

Hinrichs AS, Karolchik D, Baertsch R, Barber GP, Bejerano G, Clawson H, Diekhans M, Furey TS, Harte RA, Hsu F, et al. 2006. The UCSC Genome Browser Database: update 2006. Nucleic Acids Res 34: D590–8.

Hormozdiari F, Konkel MK, Prado-Martinez J, Chiatante G, Herraez IH, Walker JA, Nelson B, Alkan C, Sudmant PH, Huddleston J, et al. 2013. Rates and patterns of great ape retrotransposition. Proc Natl Acad Sci U S A 110: 13457–62.

Hsieh W, Chu T, Wolfinger RD, Gibson G. 2003. Mixed-model reanalysis of primate data suggests tissue and species biases in oligonucleotide-based gene expression profiles. Genetics 165: 747–57.

Jombart T. 2008. adegenet: a R package for the multivariate analysis of genetic markers. Bioinformatics 24: 1403–5.

Jombart T, Ahmed I. 2011. adegenet 1.3-1: new tools for the analysis of genome-wide SNP data. Bioinformatics 27: 3070–1.

Jonsson JJ, Foresman MD, Wilson N, Mclvor RS. 1992. Intron requirement for expression of the human purine nucleoside phosphorylase gene. Nucleic Acids Res 20: 3191–3198.

Kasen S, Ouellette R, Cohen P. 1990. Mainstreaming and postsecondary educational and employment status of a rubella cohort. Am Ann Deaf 135: 22–6.

Kashi Y, King DG. 2006. Simple sequence repeats as advantageous mutators in evolution. Trends Genet 22: 253–9.

Kelkar YD, Eckert Ka, Chiaromonte F, Makova KD. 2011. A matter of life or death: how microsatellites emerge in and vanish from the human genome. Genome Res 21: 2038–48.

Kelkar YD, Tyekucheva S, Chiaromonte F, Makova KD. 2008. The genome-wide determinants of human and chimpanzee microsatellite evolution. Genome Res 18: 30–8.

Kent WJ, Sugnet CW, Furey TS, Roskin KM, Pringle TH, Zahler AM, Haussler a. D. 2002. The Human Genome Browser at UCSC. Genome Res 12: 996–1006.

Kim S-J, Young LJ, Gonen D, Veenstra-VanderWeele J, Courchesne R, Courchesne E, Lord C, Leventhal BL, Cook EH, Insel TR. 2002. Transmission disequilibrium testing of arginine vasopressin receptor 1A (AVPR1A) polymorphisms in autism. Mol Psychiatry 7: 503–7.

King MC, Wilson AC. 1975. Evolution at two levels in humans and chimpanzees. Science 188: 107–16.

Kinsella RJ, Kähäri A, Haider S, Zamora J, Proctor G, Spudich G, Almeida-King J, Staines D, Derwent P, Kerhornou A, et al. 2011. Ensembl BioMarts: a hub for data retrieval across taxonomic space. Database (Oxford) 2011: bar030.

Lander ES, Linton LM, Birren B, Nusbaum C, Zody MC, Baldwin J, Devon K, Dewar K, Doyle M, FitzHugh W, et al. 2001. Initial sequencing and analysis of the human genome. Nature 409: 860–921.

Landry CR, Lemos B, Rifkin S a, Dickinson WJ, Hartl DL. 2007. Genetic properties influencing the evolvability of gene expression. Science 317: 118–21.

Lawrence M, Huber W, Pagès H, Aboyoun P, Carlson M, Gentleman R, Morgan MT, Carey VJ. 2013. Software for computing and annotating genomic ranges. PLoS Comput Biol 9: e1003118.

Legendre M, Pochet N, Pak T, Verstrepen KJ. 2007. Sequence-based estimation of minisatellite and microsatellite repeat variability. Genome Res 17: 1787–96.

Lehner B. 2008. Selection to minimise noise in living systems and its implications for the evolution of gene expression. Mol Syst Biol 4: 170.

Li JZ, Absher DM, Tang H, Southwick AM, Casto AM, Ramachandran S, Cann HM, Barsh GS, Feldman M, Cavalli-Sforza LL, et al. 2008. Worldwide human relationships inferred from genome-wide patterns of variation. Science 319: 1100–1104.

Li Y, Korol AB, Fahima T, Nevo E. 2004. Microsatellites within genes: structure, function, and evolution. Mol Biol Evol 21: 991–1007.

Li Y, Zheng H, Luo R, Wu H, Zhu H, Li R, Cao H, Wu B, Huang S, Shao H, et al. 2011. Structural variation in two human genomes mapped at single-nucleotide resolution by whole genome de novo assembly. Nat Biotechnol 29: 723–30.

Li Y-C, Korol AB, Fahima T, Beiles A, Nevo E. 2002. Microsatellites: genomic distribution, putative functions and mutational mechanisms: a review. Mol Ecol 11: 2453–65.

Lim KG, Kwoh CK, Hsu LY, Wirawan A. 2012. Review of tandem repeat search tools: a systematic approach to evaluating algorithmic performance. Brief Bioinform 14: 67–81.

Locke DP, Hillier LW, Warren WC, Worley KC, Nazareth LV, Muzny DM, Yang S-P, Wang Z, Chinwalla AT, Minx P, et al. 2011. Comparative and demographic analysis of orang-utan genomes. Nature 469: 529–33.

Loire E, Higuet D, Netter P, Achaz G. 2013. Evolution of coding microsatellites in primate genomes. Genome Biol Evol 5: 283–95.

Macneil LT, Walhout AJM. 2011. Gene regulatory networks and the role of robustness and stochasticity in the control of gene expression. 645–657.

Mann HB, Whitney DR. 1947. On a Test of Whether one of Two Random Variables is Stochastically Larger than the Other. Ann Math Stat 18: 50–60.

Mardis ER. 2008. The impact of next-generation sequencing technology on genetics. Trends Genet 24: 133–41.

Marques-Bonet T, Ryder Oa, Eichler EE. 2009. Sequencing primate genomes: what have we learned?. Annu Rev Genomics Hum Genet 10: 355–86.

Material SO, Web S, Press H, York N, Nw A. 2004. The ENCODE (ENCyclopedia Of DNA Elements) Project. Science 306: 636–40.

McIver LJ, Fondon JW, Skinner M a, Garner HR. 2011. Evaluation of microsatellite variation in the 1000 Genomes Project pilot studies is indicative of the quality and utility of the raw data and alignments. Genomics 97: 193–9.

McIver LJ, McCormick JF, Martin a, Fondon JW, Garner HR. 2013. Population-scale analysis of human microsatellites reveals novel sources of exonic variation. Gene 516: 328–34.

McKenna A, Hanna M, Banks E, Sivachenko A, Cibulskis K, Kernytsky A, Garimella K, Altshuler D, Gabriel S, Daly M, et al. 2010. The Genome Analysis Toolkit: a MapReduce framework for analyzing next-generation DNA sequencing data. Genome Res 20: 1297–303.

McMahon BJ, Teeling EC, Höglund J. 2014. How and why should we implement genomics into conservation?. Evol Appl 7: 999–1007.

McManus KF, Kelley JL, Song S, Veeramah KR, Woerner AE, Stevison LS, Ryder OA, Ape Genome Project G, Kidd JM, Wall JD, et al. 2015. Inference of gorilla demographic and selective history from whole-genome sequence data. Mol Biol Evol 32: 600–12.

Metzker ML. 2010. Sequencing technologies - the next generation. Nat Rev Genet 11: 31–46.

Meyer M, Kircher M, Gansauge M-T, Li H, Racimo F, Mallick S, Schraiber JG, Jay F, Prüfer K, de Filippo C, et al. 2012. A high-coverage genome sequence from an archaic Denisovan individual. Science 338: 222–6.

Molla M, Delcher A, Sunyaev S, Cantor C, Kasif S. 2009. Triplet repeat length bias and variation in the human transcriptome. Proc Natl Acad Sci U S A 106: 17095–100.

Morrison NA, Stephens AA, Osato M, Polly P, Tan TC, Yamashita N, Doecke JD, Pasco J, Fozzard N, Jones G, et al. 2012. Glutamine repeat variants in human RUNX2 associated with decreased femoral neck BMD, broadband ultrasound attenuation and target gene transactivation. PLoS One 7: e42617.

Nater A, Arora N, Greminger MP, Van Schaik CP, Singleton I, Wich SA, Fredriksson G, Perwitasari-Farajallah D, Pamungkas J, Krützen M. 2013. Marked population structure and recent migration in the critically endangered sumatran orangutan (Pongo abelii). J Hered 104: 2–13.

O’Dushlaine CT, Shields DC. 2008. Marked variation in predicted and observed variability of tandem repeat loci across the human genome. BMC Genomics 9: 175.

Paradis E, Claude J, Strimmer K. 2004. APE: Analyses of Phylogenetics and Evolution in R language. Bioinformatics 20: 289–290.

Paten B, Herrero J, Beal K, Fitzgerald S, Birney E. 2008. Enredo and Pecan: genome-wide mammalian consistency-based multiple alignment with paralogs. Genome Res 18: 1814–28.

Payseur B a, Jing P, Haasl RJ. 2011. A genomic portrait of human microsatellite variation. Mol Biol Evol 28: 303–12.

Pemberton TJ, DeGiorgio M, Rosenberg NA. 2013. Population structure in a comprehensive genomic data set on human microsatellite variation. G3 (Bethesda) 3: 891–907.

Pemberton TJ, Sandefur CI, Jakobsson M, Rosenberg N a. 2009. Sequence determinants of human microsatellite variability. BMC Genomics 10: 612.

Pilpel Y. 2011. Noise in biological systems: pros, cons, and mechanisms of control. eds. J.I. Castrillo and S.G. Oliver. Methods Mol Biol 759: 407–25.

Prado-Martinez J, Sudmant PH, Kidd JM, Li H, Kelley JL, Lorente-Galdos B, Veeramah KR, Woerner AE, O’Connor TD, Santpere G, et al. 2013. Great ape genetic diversity and population history. Nature 499: 471–5.

Press MO, Carlson KD, Queitsch C. 2014. The overdue promise of short tandem repeat variation for heritability. Trends Genet. 0–27.

Pritchard JK, Stephens M, Donnelly P. 2000. Inference of population structure using multilocus genotype data. Genetics 155: 945–59.

Prüfer K, Racimo F, Patterson N, Jay F, Sankararaman S, Sawyer S, Heinze A, Renaud G, Sudmant PH, de Filippo C, et al. 2014. The complete genome sequence of a Neanderthal from the Altai Mountains. Nature 505: 43–9.

Raveh-Sadka T, Levo M, Shabi U, Shany B, Keren L, Lotan-Pompan M, Zeevi D, Sharon E, Weinberger A, Segal E. 2012. Manipulating nucleosome disfavoring sequences allows fine-tune regulation of gene expression in yeast. Nat Genet 44: 743–750.

Reinartz GE, Karron JD, Phillips RB, Weber JL. 2000. Patterns of microsatellite polymorphism in the range-restricted bonobo (Pan paniscus): considerations for interspecific comparison with chimpanzees (P. troglodytes). Mol Ecol 9: 315–328.

Rockman MV, Wray GA. 2002. Abundant raw material for cis-regulatory evolution in humans. Mol Biol Evol 19: 1991–2004.

Rohrer J, Conley ME. 1998. Transcriptional regulatory elements within the first intron of Bruton’s tyrosine kinase. Blood 91: 214–21.

Rosenberg NA, Pritchard JK, Weber JL, Cann HM, Kidd KK, Zhivotovsky LA, Feldman MW. 2002. Genetic structure of human populations. Science 298: 2381–5.

Sabo PJ, Kuehn MS, Thurman R, Johnson BE, Johnson EM, Cao H, Yu M, Rosenzweig E, Goldy J, Haydock A, et al. 2006. Genome-scale mapping of DNase I sensitivity in vivo using tiling DNA microarrays. Nat Methods 3: 511–518.

Sawaya S, Bagshaw A, Buschiazzo E, Kumar P, Chowdhury S, Black MA, Gemmell N. 2013. Microsatellite tandem repeats are abundant in human promoters and are associated with regulatory elements. PLoS One 8: e54710.

Saxonov S, Berg P, Brutlag DL. 2006. A genome-wide analysis of CpG dinucleotides in the human genome distinguishes two distinct classes of promoters. Proc Natl Acad Sci U S A 103: 1412–1417.

Scally A, Dutheil JY, Hillier LW, Jordan GE, Goodhead I, Herrero J, Hobolth A, Lappalainen T, Mailund T, Marques-Bonet T, et al. 2012. Insights into hominid evolution from the gorilla genome sequence. Nature 483: 169–75.

Scally A, Yngvadottir B, Xue Y, Ayub Q, Durbin R, Tyler-Smith C. 2013. A genome-wide survey of genetic variation in gorillas using reduced representation sequencing. PLoS One 8: e65066.

Schuster SC, Miller W, Ratan A, Tomsho LP, Giardine B, Kasson LR, Harris RS, Petersen DC, Zhao F, Qi J, et al. 2010. Complete Khoisan and Bantu genomes from southern Africa. Nature 463: 943–7.

Sequencing TC, Consortium A. 2005. Initial sequence of the chimpanzee genome and comparison with the human genome. Nature 437: 69–87.

Slatkin M. 1995. A measure of population subdivision based on microsatellite allele frequencies. Genetics 139: 457–62.

Somel M, Creely H, Franz H, Mueller U, Lachmann M, Khaitovich P, Pääbo S. 2008. Human and chimpanzee gene expression differences replicated in mice fed different diets. PLoS One 3: e1504.

Spitz F, Furlong EEM. 2012. Transcription factors: from enhancer binding to developmental control. Nat Rev Genet 13: 613–26.

Stranger BE, Forrest MS, Clark AG, Minichiello MJ, Deutsch S, Lyle R, Hunt S, Kahl B, Antonarakis SE, Tavaré S, et al. 2005. Genome-wide associations of gene expression variation in humans. PLoS Genet 1: e78.

Stranger BE, Forrest MS, Dunning M, Ingle CE, Beazley C, Thorne N, Redon R, Bird CP, de Grassi A, Lee C, et al. 2007. Relative impact of nucleotide and copy number variation on gene expression phenotypes. Science 315: 848–53.

Sudmant PH, Huddleston J, Catacchio CR, Malig M, Hillier LW, Baker C, Mohajeri K, Kondova I, Bontrop RE, Persengiev S, et al. 2013. Evolution and diversity of copy number variation in the great ape lineage. Genome Res 23: 1373–82.

Sun JX, Helgason A, Masson G, Ebenesersdóttir SS, Li H, Mallick S, Gnerre S, Patterson N, Kong A, Reich D, et al. 2012. A direct characterization of human mutation based on microsatellites. Nat Genet 44: 1161–5.

Tirosh I, Barkai N, Verstrepen KJ. 2009. Promoter architecture and the evolvability of gene expression. J Biol 8: 95.

Tirosh I, Weinberger A, Carmi M, Barkai N. 2006. A genetic signature of interspecies variations in gene expression. Nat Genet 38: 830–4.

Tishkoff SA, Reed FA, Friedlaender FR, Ehret C, Ranciaro A, Froment A, Hirbo JB, Awomoyi AA, Bodo J-M, Doumbo O, et al. 2009. The genetic structure and history of Africans and African Americans. Science 324: 1035–44.

Vallender EJ. 2011. Expanding whole exome resequencing into non-human primates. Genome Biol 12: R87.

Verstrepen KJ, Jansen A, Lewitter F, Fink GR. 2005. Intragenic tandem repeats generate functional variability 37: 986–990.

Vinces MD, Legendre M, Caldara M, Hagihara M, Verstrepen KJ. 2009. Unstable tandem repeats in promoters confer transcriptional evolvability. Science 324: 1213–6.

Waal De FBM. 2009. The Age of Empathy. Harmony Books, New York, NY.

Warren KS, Nijmian IJ, Lenstra JA, Swan RA, Heriyanto, den Boer M. 2000. Microsatellite DNA variation in Bornean orangutans (Pongo pygmaeus). J Med Primatol 29: 57–62.

Weber JL, Wong C. 1993. Mutation of human short tandem repeats. Hum Mol Genet 2: 1123–8.

Webster MT. 2003. Compositional Evolution of Noncoding DNA in the Human and Chimpanzee Genomes. Mol Biol Evol 20: 278–286.

Webster MT, Smith NGC, Ellegren H. 2002. Microsatellite evolution inferred from human-chimpanzee genomic sequence alignments. Proc Natl Acad Sci U S A 99: 8748–53.

Wegmann D, Excoffier L. 2010. Bayesian inference of the demographic history of chimpanzees. Mol Biol Evol 27: 1425–35.

Wickham H. 2009. ggplot2. Springer New York, New York, NY.

Willems T, Gymrek M, Highnam G, Mittelman D, Erlich Y. 2014. The landscape of human STR variation. Genome Res 24: 1894–904.

Wray G a, Hahn MW, Abouheif E, Balhoff JP, Pizer M, Rockman MV, Romano La. 2003. The evolution of transcriptional regulation in eukaryotes. Mol Biol Evol 20: 1377–419.

Wren JD, Forgacs E, Fondon JW, Pertsemlidis A, Cheng SY, Gallardo T, Williams RS, Shohet RV, Minna JD, Garner HR. 2000. Repeat polymorphisms within gene regions: phenotypic and evolutionary implications. Am J Hum Genet 67: 345–56.

Yoon OK, Hsu TY, Im JH, Brem RB. 2012. Genetics and regulatory impact of alternative polyadenylation in human B-lymphoblastoid cells. PLoS Genet 8: e1002882.

Zhang Y, Morin PA, Ryder OA, Zhang Y. 2001. A set of human tri- and tetra-nucleotide microsatellite loci useful for population analyses in gorillas ( Gorilla gorilla gorilla) and orangutans (Pongo pygmaeus). Conserv Genet 9700: 391–395.

